# Tolerance Mechanisms in Polysaccharide Biosynthesis: Implications for Undecaprenol Phosphate Recycling in *Escherichia coli* and *Shigella flexneri*

**DOI:** 10.1101/2024.09.26.615265

**Authors:** Jilong Qin, Yaoqin Hong, Renato Morona, Makrina Totsika

## Abstract

Bacterial polysaccharide synthesis is catalysed on the universal lipid carrier, undecaprenol phosphate (UndP). The cellular UndP pool is shared by other polysaccharide synthesis pathways and in peptidoglycan (PG) biogenesis. Disruptions in cytosolic polysaccharide synthesis steps are detrimental to bacterial survival due to affecting UndP recycling. In contrast, bacteria can survive disruptions in the periplasmic steps, suggesting a tolerance mechanism to mitigate UndP sequestration. Here we investigated tolerance mechanisms to disruptions of polymerases that are involved in UndP-releasing steps in two related polysaccharide synthesis pathways: the enterobacterial common antigen (ECA) and the O antigen (OAg), in *Escherichia coli* and *Shigella flexneri*. Our study reveals that polysaccharide polymerisation is crucial for efficient UndP recycling. In *E. coli* K-12, cell survival upon disruptions in OAg polymerase is dependent on a functional ECA synthesis pathway and vice versa. This is because disruptions in OAg synthesis leads to the redirection of the shared lipid-linked sugar substrate UndPP-GlcNAc towards increased ECA production. Conversely, in *S. flexneri*, the OAg polymerase is essential due to its limited ECA production, which inadequately redirects UndP flow to support cell survival. We propose a model whereby sharing the initial sugar intermediate UndPP-GlcNAc between the ECA and OAg synthesis pathways allows UndP to be redirected towards ECA production, mitigating sequestration issues caused by disruptions in the OAg pathway. These findings suggest an evolutionary buffering mechanism that enhances bacterial survival when UndP sequestration occurs due to stalled polysaccharide biosynthesis, which may allow polysaccharide diversity in the species to increase over time.

**Importance:** Enzymes involved in bacterial polysaccharide biosynthesis have substrate specificity to ensure the correct polysaccharide is produced at the appropriate place and time. However, this specificity poses a challenge for the diversification of polysaccharide structure and hence function, as the acquisition of a novel oligosaccharide RU would likely disrupt the synthesis pathways, leading to sequestration of the essential universal lipid carrier UndP and ultimately cause cell death. We investigated how cells tolerate disruptions in polysaccharide synthesis pathways and provide evidence that suggests that sharing a common substrate between the synthesis pathways of two common enteric bacterial surface polysaccharides (ECA and OAg), can redirect the flow of UndP. Our study provides insights into the mechanism of how bacteria alleviate sequestration issues, thereby enhancing cell survival which may allow them additional capacity for polysaccharide diversification.

## Introduction

Surface polysaccharides confer enteric bacteria with resistance to host antimicrobials (1, 2) and ability to colonise various host niches (3). The O antigen (OAg) polysaccharide made of oligosaccharide repeating units (RUs) represents an important virulence factor for Gram-negative bacteria contributing to host colonisation (4) and virulence modulation (5). The structure of OAg is under strong selection by host immunity and host niche-residing bacteriophages, giving rise to over 180 diverse OAg structures characterised to date in *Escherichia coli* (including *Shigella* strains) (6), which form the molecular basis of the O-typing scheme (7).

The synthesis and assembly of OAg rely on the universal lipid carrier undecaprenol phosphate (UndP or C_55_), which is shared with the biosynthesis of other polysaccharides, such as enterobacterial common antigen (ECA) and peptidoglycan (PG) in the cell (Fig 1A). However, cells only produce a finite amount of UndP (<1% of total membrane lipids (8)) through *de novo* synthesis (Fig 1A), and the demand for UndP during synthesis of different polysaccharides is critically ensured through efficient recycling (Fig 1A). In the biosynthesis cycle of OAg and ECA for most *E. coli* and *S. flexneri*, UndP is engaged by the initial transferase (IT), *N*-acetylglucosamine-1-phosphate transferase WecA, to form UndPP-GlcNAc in an enzymatically reversible manner (9) (Fig 1B). UndP is then committed to either the OAg or ECA synthesis cycle in the subsequent step catalysed by the second glycosyltransferase (GT), WbbL (for OAg in *E. coli* K-12 (10)) or WecG (for ECA), and remains occupied by RUs during their assembly (Fig 1B), including the subsequent membrane-flipping step by flippases WzxB and WzxE, respectively (Fig 1A). UndPP is then released in the periplasm (PP) from UndPP-OAg (by OAg polymerase WzyB and ligase WaaL), and UndPP-ECA (by ECA polymerase WzyE and ligase WaaL) (Fig 1A). While for PG synthesis, UndP is engaged after RU assembly by MraY, translocated by MurJ, and released by penicillin binding proteins (PBPs) and proteins belonging to the SEDS (shape, elongation, division, and sporulation) family (Fig 1A). The released UndPP is further dephosphorylated into UndP by pyrophosphatases and flipped back into the cytosolic leaflet for subsequent rounds of synthesis (Fig 1A).

**Fig 1.**
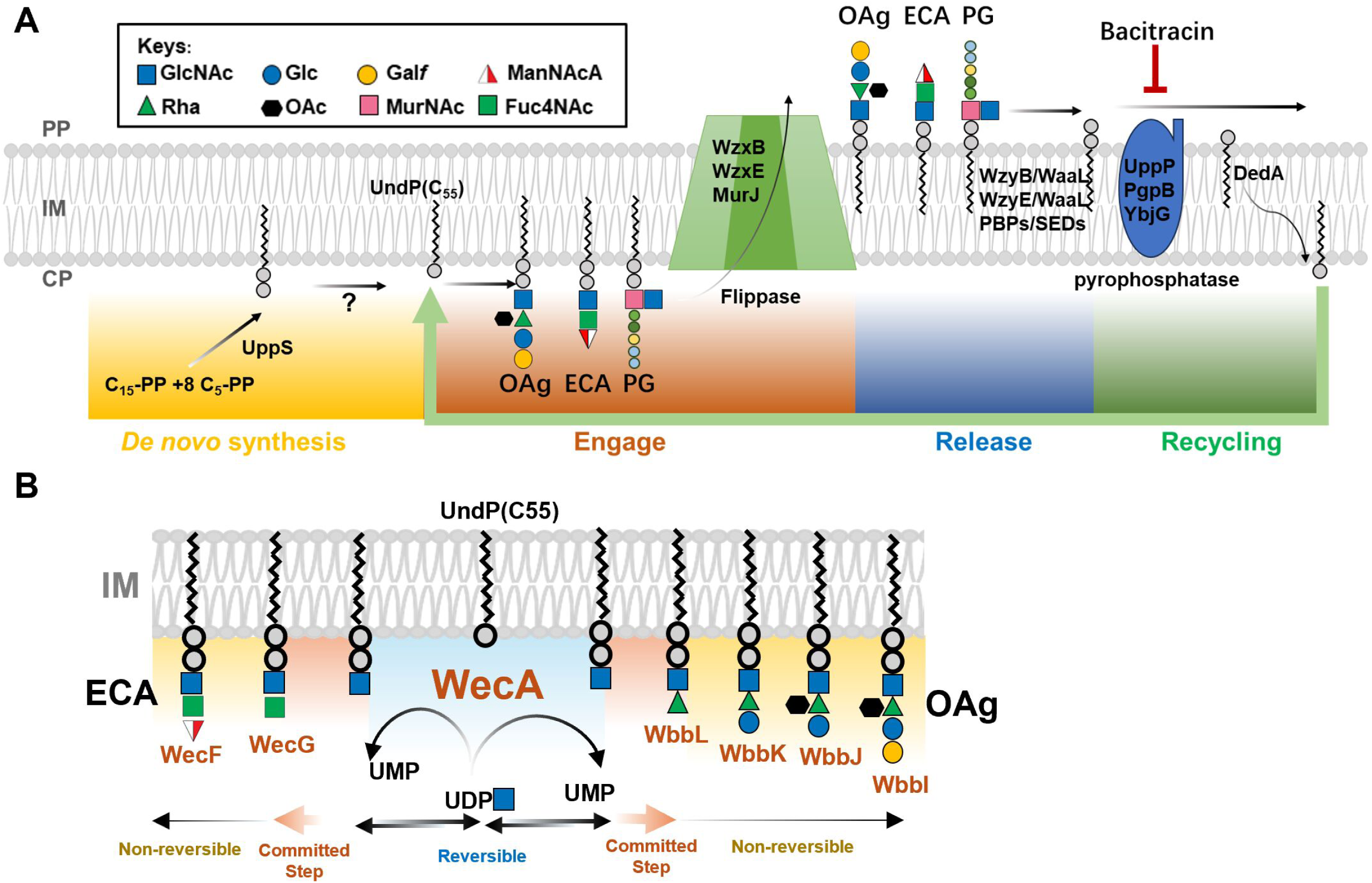
Biogenesis of bacterial polysaccharides on the universal lipid carrier UndP. **A)** Schematic representation of UndP *de novo* synthesis, glycan RU engagement, release and recycling during OAg, ECA and PG biosynthesis in *E. coli*. PP, periplasm; IM inner membrane; CP, cytoplasm. **B)** Schematic representation of the biogenesis of ECA and OAg RUs on the UndP lipid carrier with the shared initial saccharide GlcNAc. GTs responsible for saccharide assembly are shown in orange.

The enzymes collectively responsible for the biosynthesis and assembly of different polysaccharides recognise the substrates with a common feature, UndPP-linked glycans. To ensure that the different polysaccharides are synthesised and assembled correctly, synthesis pathways have specificities towards glycan moieties on the lipid carrier in three different processes: step-wised assembly of UndPP-RUs through glycosyltransferases (GT), IM translocation of UndPP-RUs through flippases, and polymerisation of RUs through polymerases. The specificity of GTs is also towards acceptors (UndPP-glycans), and disruptions in GTs will abolish or stall the RU assembly on UndPP (11). Flippase specificity ensures that only the complete UndPP-RUs are efficiently translocated across the IM, leaving any incomplete RUs in the cytosol (12). Polymerases are specific towards UndPP-RU that only efficiently polymerise the correct RUs (13). Therefore, genes coding for OAg GTs, flippases and polymerases have high sequence diversity and are often specific to an individual OAg gene cluster, a feature that has been be exploited as a genotyping method for rapid strain identification and clinical detection (14).

Although enzyme specificities are crucial to the quality control of polysaccharide biosynthesis, they may represent a limitation in the evolutionary diversification process for their polysaccharide RU structures for serotype switching. We have previously shown that expression of the mono-rhamnosyltransferase WbbL in *S. flexneri* that is defective in di-rhamnosyltransferase RfbG resulted in cell lysis (11). This is because the incomplete OAg RU intermediate UndPP-GlcNAc-Rha catalysed by WbbL is not recognised neither by the last GT RfbF nor the flippase WzxB in *S. flexneri*, causing sequestration of UndP in OAg biosynthesis, thereby limiting the availability of UndP for PG synthesis. This observation suggested that the evolution of a new OAg RU type (serotype switching) comes with a risk of deleterious effects, in that the newly incorporated functional GT may stall the existing OAg synthesis due to specificities of polysaccharide enzymes, leading to UndP sequestration and cell death. Therefore, understanding bacterial tolerance to UndP sequestration, particularly in the context of disruptions to synthesis pathways, can provide insights into the evolution of polysaccharide diversification, a biological process that is widely common in bacteria.

Here we studied the survival and tolerance to UndP sequestration in *E. coli* and *S. flexneri* mutants defective in UndPP releasing steps during both OAg and ECA synthesis. We showed that *E. coli* and *S. flexneri* have different tolerance to the disruption of OAg polymerases. The high tolerance to OAg polymerase disruption in *E. coli* was due to increased redirection of the substrate UndPP-GlcNAc towards ECA synthesis, whereas the low tolerance to OAg polymerase disruption in *S. flexneri* was due to its low ECA synthesis capacity. Our data suggest a buffering mechanism, where through sharing the initial substrate UndPP-GlcNAc between OAg and ECA synthesis, *E. coli* can alleviate UndP sequestration stress during OAg synthesis by redirecting UndP flow to the ECA synthesis cycle and ensure increased cell survival.

## Results

### The OAg polymerase WzyB contributes to rapid UndP recycling

Biosynthesis of OAg occupies a pool of UndP (15). Consequently, restoring OAg synthesis in *E. coli* K-12 strain MG1655-S (16) increased sensitivity towards bacitracin, an antibiotic targeting the pyrophosphatases for UndP recycling (Fig 1A), without affecting cell viability (Fig 2A). Limiting the level of UndP by sequestering it in OAg or ECA synthesis has a negative impact on cell survival as it limits the synthesis of essential cell wall component PG. In *E. coli* K-12 strain MG1655, switching on OAg production by restoring the committed-step GT WbbL in a flippase *wzxB* mutant background inhibits cell growth (Fig 2B). Disruption of *wzxB* has been shown to sequester UndP in UndPP-OAg intermediates (15) with documented cell shape deformities (17), a hallmark of PG synthesis defects. These results collectively confirmed that sequestration of UndP in the UndPP-linked OAg RU intermediates in the cytosolic face of IM is lethal. In contrast, in the WbbL-complemented MG1655 (Fig 2C), disruption of the OAg ligase WaaL, known to sequester UndP in the UndPP-OAg intermediates in the periplasmic leaflet (15, 16), is not lethal. We reasoned that the polymerisation reaction by WzyB in the absence of WaaL releases enough UndPP to be recycled for PG synthesis (Fig 2D), and therefore only the disruption of both WaaL and WzyB is lethal. Indeed, while disruption of either WaaL or WzyB has no impact on cell growth, disruption of both WzyB and WaaL inhibited cell growth (Fig 2C). These results suggested that the OAg polymerase WzyB contributes to rapid UndP recycling. Our findings are in line with a previous report showing that disruption of *wzyB* accumulated a high level of UndPP-OAg intermediates (15).

**Fig 2.**
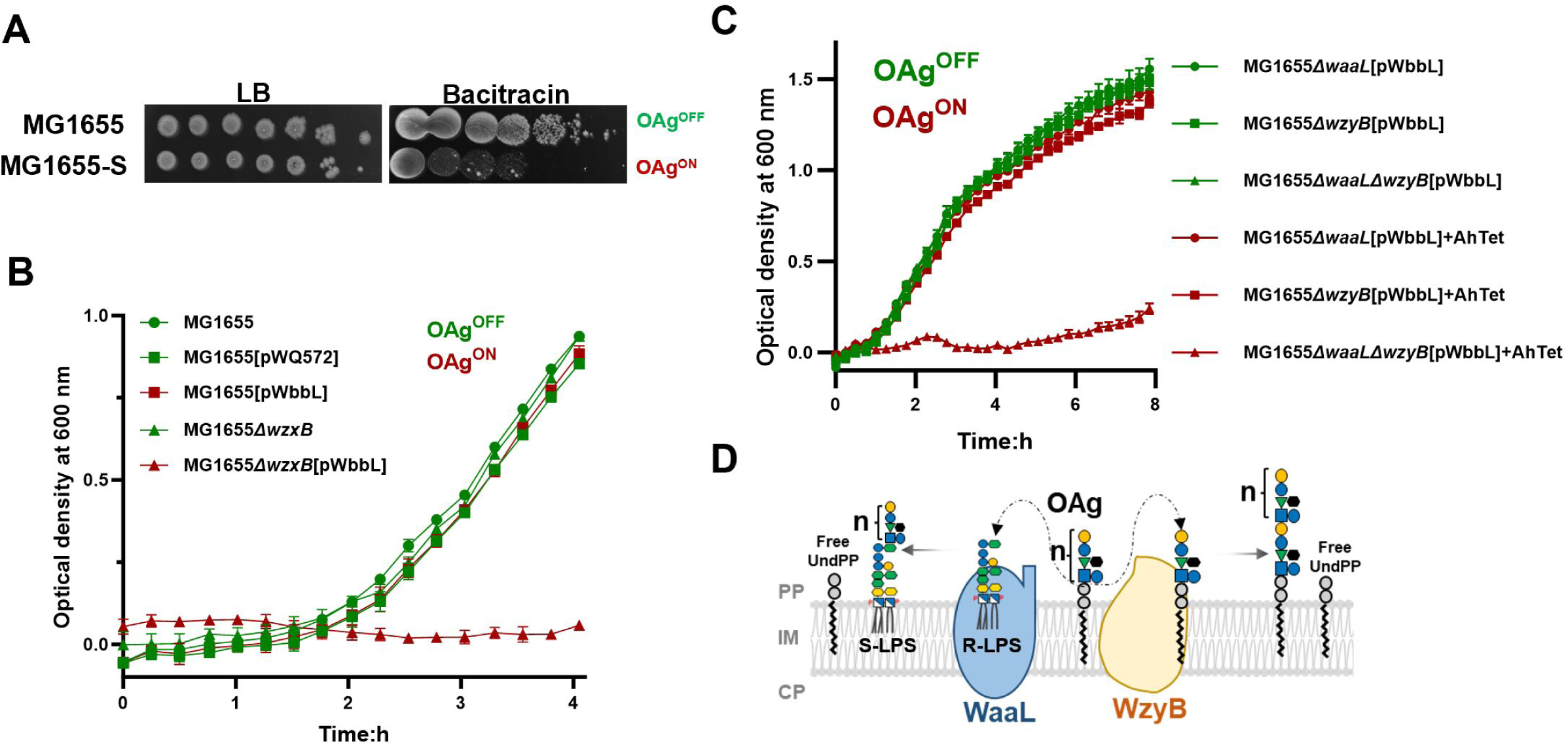
Synthetic lethality of Δ*waaL*Δ*wzyB* double knockouts due to complete stalling of the UndPP-OAg intermediates. **A)** Bacitracin sensitivity assay for *E. coli* MG1655 and MG1655-S. Overnight bacterial cultures were 10-folded diluted and spotted on LB agar plates supplemented without or with 1 mg/ml bacitracin. **B-C)** Growth curves of *E. coli* K-12 strains harbouring plasmids without or with *wbbL* cultured in LB media supplemented with or without anhydrotetracycline (AnTet). The status of OAg production is labelled as OAg^ON^ in red, or OAg^OFF^ in green. **D)** Schematic representation of enzymatic reactions catalysed by WaaL and WzyB in releasing UndPP from UndPP-OAg RUs. n=17-21 in *E. coli* K-12 for O16-OAg.

### Disruption of wzyB in S. flexneri is lethal

Interestingly, the gene encoding OAg polymerase WzyB in *S. flexneri* was reported to lack transposon insertions in a dense transposon insertion library (18), strongly indicating that it is essential. Here, we found that attempting to construct a direct *wzyB* deletion in *S. flexneri* resulted in a few small and slow-growing colonies (Fig 3A). By diagnostic PCR it was shown that the putative *wzyB* mutant contains a *wzyB* duplication, which likely acts as a suppressor mutation (Fig 3B). Since WzyB contributes to rapid UndP recycling, we suspected that disruption of *wzyB* may cause UndP sequestration in *S. flexneri*, impacting its growth. We therefore made the *wzyB* deletion in an IT Δ*wecA* mutant background (which is disrupted for UndP engagement in both ECA and OAg synthesis) to avoid potential UndP sequestration and confirmed this to be the case by successful deletion of *wzyB* in this background (Fig 3A-B). Strikingly, complementation of WecA expression in the *ΔwecAΔwzyB* double mutant resulted in cell lysis with the release of cellular DNA into culture supernatant (Fig 3C). These results suggest that WzyB is essential in *S. flexneri,* in that the deletion of *wzyB* causes UndP sequestration leading to cell death.

**Fig 3.**
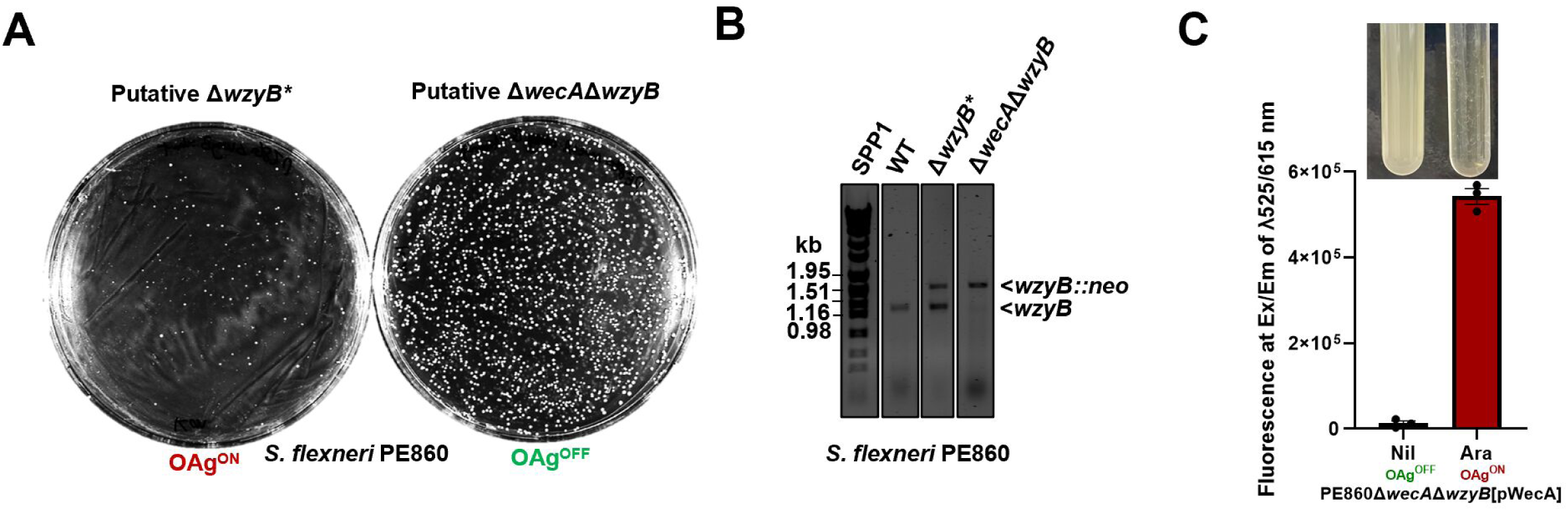
WzyB is essential in *S. flexneri.* **A)** Colony number and morphology of putative *wzyB* inactivation mutants in either the background of WT or *wecA*-null *S. flexneri*. **B)** Diagnostic PCR for *wzyB* inactivation with primers targeting coding sequences of WzyB. Successful insertions of *neo* in *wzyB* is indicated as *wzyB::neo*. **C)** Amount of cellular DNA detected by ethidium bromide (EtBr) in *S. flexneri* Δ*wecA*Δ*wzyB* culture supernatant released upon pWecA induction. Cell lysis was imaged as loss of turbidity in culture media. The status of OAg production is labelled as either OAg^ON^ in Red or OAg^OFF^ in Green.

### Redirection of UndPP-GlcNAc between ECA and OAg to mitigate UndP sequestration

Besides the OAg polymerase WzyB being essential in *S. flexneri*, the ECA polymerase WzyE was also found essential in *E. coli* K-12 strains lacking OAg, while not in *E. coli* strains producing OAg (ST131) (19). Moreover, we could delete *wzyE* in the OAg restored *E. coli* K-12 strain MG1655-S. We also successfully deleted *wzyE* in MG1655-S*ΔwecA* (disruption in engaging UndP for both OAg and ECA synthesis). In this strain, switching on both ECA and OAg production by inducing WecA in a *ΔwecAΔwzyE* double mutant revealed no growth defects, suggesting that when OAg is produced, WzyE is not essential (Fig 4A). We reasoned that this is because, in most *E. coli* strains, OAg shares the same first sugar *N*-acetylglucosamine (GlcNAc) with ECA, allowing the common substrate UndPP-GlcNAc to be redirected into making OAg (Fig 1B) when UndP is sequestered in a *wzyE* mutant. This prompted us to examine the essentiality of the OAg polymerase WzyB in the OAg-restored *E. coli* K-12 strain with ECA synthesis inactivated (MG1655-S*ΔwecAΔwecGΔwzyB*). Intriguingly, when ECA synthesis is inactivated through the disruption of committed step catalysed by WecG, induction of WecA in MG1655-S*ΔwecAΔwecGΔwzyB* resulted in cell lysis (Figure 4B-C), suggesting that the OAg polymerase WzyB is essential when ECA synthesis is inactivated. Together, these results led us to propose a model that by sharing a common initiating sugar GlcNAc between ECA and OAg, bacteria could gain increased tolerance to UndP sequestration through redirecting their common substrate UndPP-GlcNAc into remaining functional synthesis pathways to ensure rapid UndP recycling.

**Fig 4.**
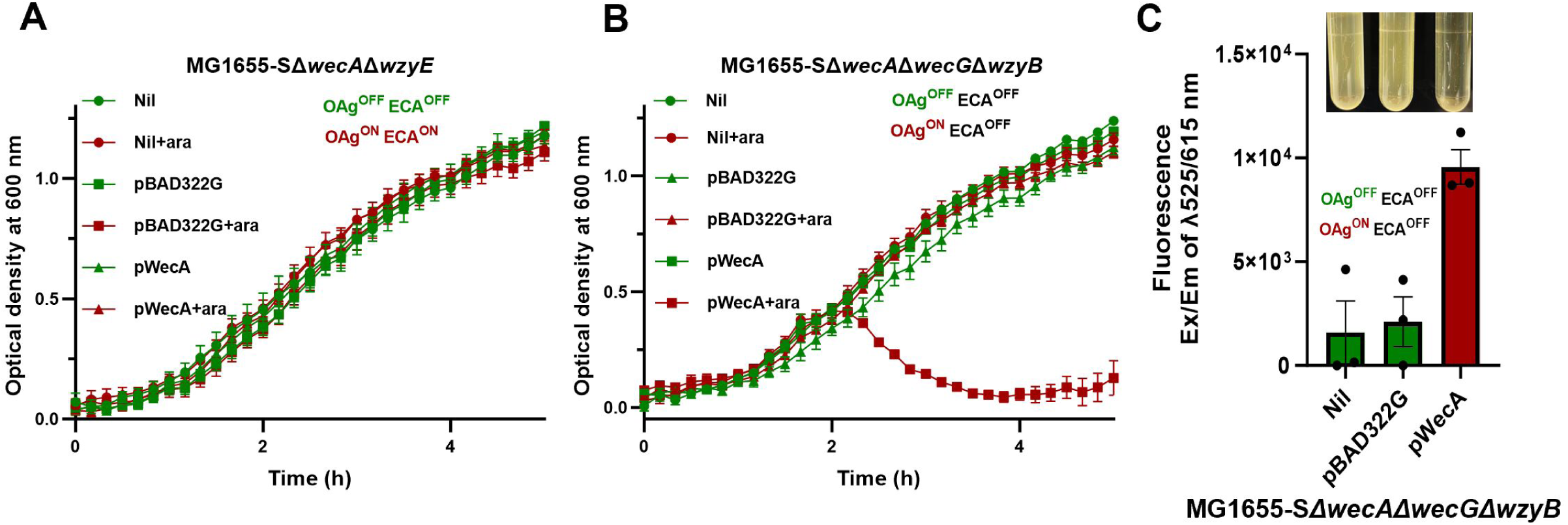
Interdependence of *wzy* essentiality between OAg and ECA synthesis pathways. A&B) Growth curves of indicated *E. coli* K-12 strains harbouring plasmids either without or with *wecA* cultured in LB media supplemented either with or without arabinose (ara). **C)** Release of cellular DNA of MG1655-SΔ*wecA*Δ*wecG*Δ*wzyB* in culture supernatant upon pWecA induction detected by ethidium bromide (EtBr). Cell lysis was imaged as loss of turbidity in culture media. The status of OAg production is labelled as either OAg^ON^ in Red or OAg^OFF^ in Green.

### Low tolerance to UndP sequestration in S. flexneri due to limited ECA production

OAg in both WbbL-restored *E. coli* K-12 and *S. flexneri* initiates with GlcNAc, yet showed different tolerance to cell survival due to UndP sequestration upon *wzyB* disruption (Fig 2C&3C). This prompted us to examine ECA production in both strains. We first confirmed that disruptions of both *wecG* and *wzyE* abolished the detection of ECA in both MG1655 and MG1655-S with anti-ECA antibodies, showing that the antibody is specific. Consistent with our model raised above, *E. coli* K-12 lacking OAg (MG1655) produced a high level of ECA in comparison to its OAg restored strain MG1655-S (Fig 5A). In contrast, in *S. flexneri* strain 2457T, inactivation of OAg biosynthesis by disruption of *rmlD*, involved in the synthesis of the precursor for the second sugar, L-Rhamnose within the OAg RU, only marginally increased the ECA production level (Fig 5A). Mature ECA is predominantly located on the bacterial cell surface, and we therefore examined ECA surface production in both *E. coli* and *S. flexneri*. While the production of OAg on the cell surface masks ECA detection by anti-ECA antibodies for both *E. coli* K-12 MG1655-S and *S. flexneri* 2457T (Fig 5B), inactivation of OAg production unexpectedly revealed that surface ECA could only be detected on approximately 30% of the *S. flexneri* Δ*rmlD* population, in comparison to 95% for *E. coli* K-12 (Fig 5B-C). These results in part may explain the different tolerance to *wzyB* disruption between *E. coli* K-12 and *S. flexneri* (Fig 2C&3C), in that the capacity to make ECA in the *S. flexneri* Δ*rmlD* cell population is limited (30%) in comparison to *E. coli* MG1655 cell population (95%), leading to inadequate redirection of UndPP-GlcNAc into making ECA when *wzyB* is disrupted for the majority of the cell population (∼70%). This limited ECA production in *S. flexneri* when OAg is disrupted renders it more susceptible to UndP sequestration, and ultimately cell death when WzyB is disrupted.

**Fig 5.**
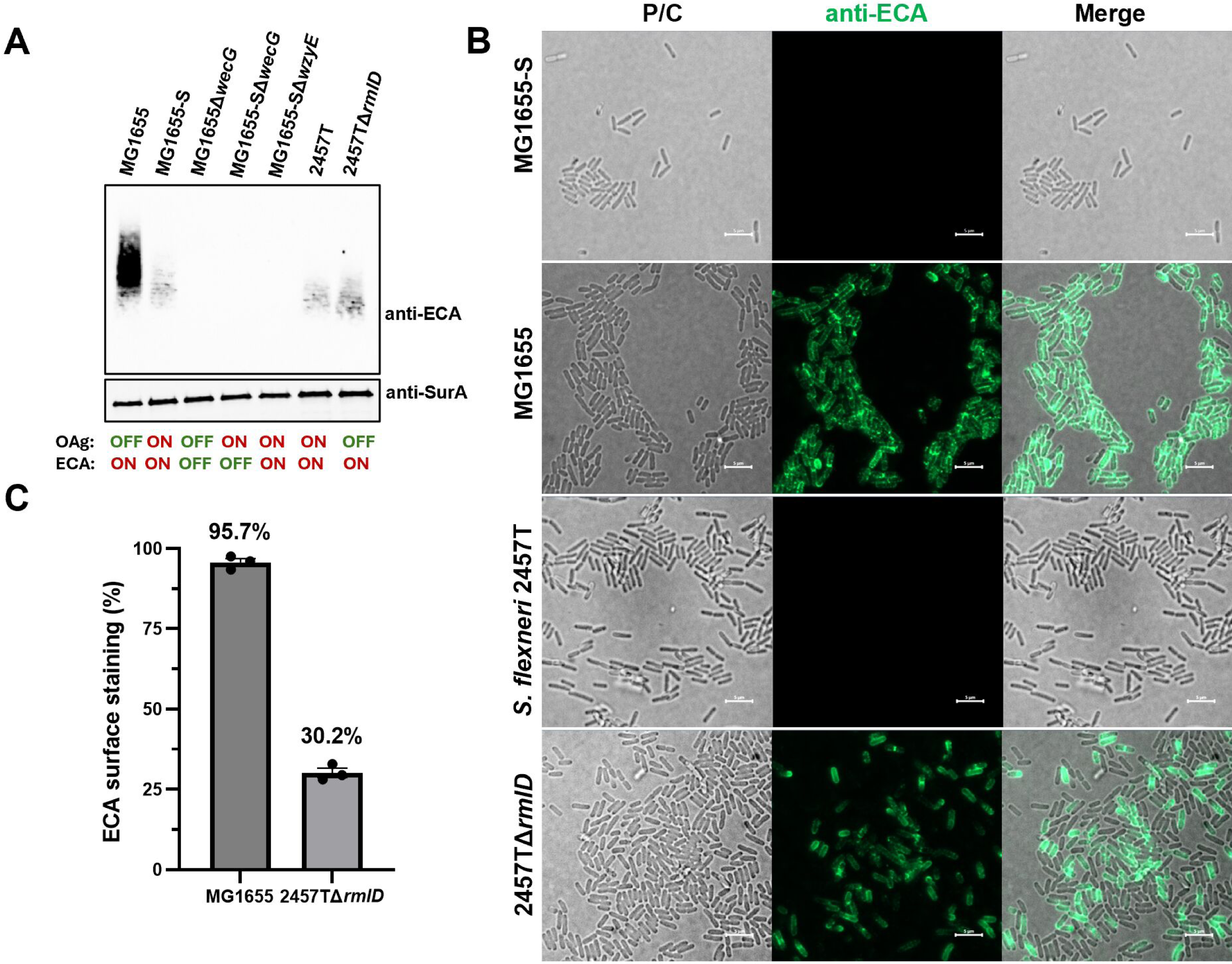
Different ECA biosynthesis levels revealed between *E. coli* K-12 and *S. flexneri* upon OAg inactivation. **A)** Western immunoblots of ECA of whole bacterial lysis samples from indicated bacterial strains. Detection of periplasmic marker SurA was used as a loading control. The status of OAg and ECA biogenesis for each strain is marked as OFF and ON. **B)** Surface ECA immunodetection via Epifluorescence microscopy. Scale bar shown as 5 μm. **C)** quantification of ECA stained bacterial population in percentages for indicated bacterial strains from three micrographs.

## Discussion

Polysaccharide synthesis requires a cellular pool of universal lipid carrier UndP. Indeed, the production of OAg although had no observable impact on growth, was shown to occupy a low level of UndP in the UndPP-RU intermediates (15), and this was also supported here by the increased sensitivity of OAg-producing *E. col*i K-12 to bacitracin, an antibiotic that limits the availability of functional UndP. When the OAg synthesis pathway is disrupted beyond the committed steps, UndP was shown to be sequestered in UndPP-RU intermediates at a high level (15), severely affecting bacterial survival (11) due to affecting the PG synthesis (17). Therefore, rapid assembly of polysaccharide RU on UndP carrier is favoured as it would lower the demand on the UndP at any given time. The rapid polysaccharide assembly could likely be achieved via substrate channelling where GTs could form a complex to fast-process UndPP-RU intermediates towards releasing. However, evidence of the formation of protein complexes by GTs encoded in the *rfb* region is currently lacking and remains to be confirmed. The other possible strategy is through GT gene arrangement in the *rfb* region. In at least 39% of OAg gene clusters (*rfb* region, including K-12 O16 *rfb*), the genes for GTs are organised in a reverse order relative to their functional steps in the biochemical process, in that the GT responsible for the committed step is located at the most downstream position (6). Consequently, this may generate an enzyme concentration gradient with the lowest level of GTs responsible for the committed steps and highest level of GTs responsible for last steps of UndPP-RU assembly, ensuring a low level of commitment of UndP to the synthesis pathways and an accelerated assembly of the complete UndPP-RU process thereafter for a rapid release of UndPP back into the cellular pool.

Disruption of OAg ligase *waaL* was shown previously to sequester high levels of UndPP-RU in OAg-producing bacterial strains (15). However, an OAg-producing *E. coli* K-12 with *waaL* deletion had no growth impact here and was confirmed to lack suppressor mutations by whole genome sequencing. We have revealed here that the polymerase WzyB which consolidates the RUs into polymeric forms and therefore rapidly releases UndPP in the absence of WaaL in the periplasm is crucial for cell viability. In a previous study (15), the level of C^14^ labelled OAg RUs on the isolated UndP pool was quantified in both *wzyB* and *waaL* mutants, with higher signal level of UndPP-linked OAg RU detected in the *wzyB* mutant. In the *wzyB* mutant, the stoichiometric relationship between OAg RU and UndP is 1:1, whereas it is (17–21):1 in the *waaL* mutant. Therefore, this suggests that a *wzyB* mutant sequestered a substantially higher level of UndP than a *waaL* mutant, suggesting that WzyB contributes to the rapid release of UndP from the OAg synthesis pathway. Supporting this tenet, we have also found that in *S. flexneri*, deletion of *wzyB* is lethal. However, two *wzyB* mutants in *S. flexneri* have been reported previously. One of the only two reported *S. flexneri wzyB* mutants was acquired through Sf6c phage selection (*S. flexneri* Y OAg as its primary receptor) (20). However the mutant could not be fully complemented to confer resistance to Colicin E2 to WT level (21), a toxin whose entry is blocked by polymerised OAg (22). Moreover, this mutant was later also used to derive a Δ*wzyB*Δ*waaL* double mutant in *S. flexneri* with no reported impact on growth (23), whereas it was shown here that deletions of both *wzyB* and *waaL* is lethal due to completely stalled UndPP-OAg intermediates. The other previously reported *S. flexneri wzyB* mutant (24) was acquired through direct allelic exchange mutagenesis after numerous attempts in the laboratory. The colony morphology of those putative mutants on the mutagenesis selection plate was similar as reported here, being overall small and heterogeneous in size. Like the other *S. flexneri wzyB* mutant that resulted from phage selection, this *wzyB* mutant could not be complemented to produce smooth-LPS (S-LPS) to WT level (24). Therefore, we believe that the two previously reported *wzyB* mutants likely contain suppressor mutations that have tuned down the committed step of OAg synthesis to reduce UndP sequestration for fitness, similar to suppressor mutations in *rml* genes reported previously in an OAg producing *E. coli* K-12, making OAg at a reduced level (16). Together, these results caution future investigations on the impacts of potential UndP sequestration when disrupting genes responsible for polysaccharide synthesis, highlighting the necessity of conducting genetic studies on these genes using an expression controlled system at IT or committed GT step (11). Indeed, genetic instability associated with polysaccharide biosynthetic defects was well recognised by Nikaido and colleagues since 1969, where studies in the galactose-initiated pathway would control synthesis with exogenously supplied D-galactose (25).

Contrasting to the essentiality of *wzyB* reported here in *S. flexneri*, a *wzyB* mutant was found to have no impact on growth in an OAg-producing *E. coli* K-12, and was reported previously with no suppressor mutations (16), hence being genetically stable. Both OAg of *S. flexneri* and *E. coli* K-12 are initiated with GlcNAc, which is the same initiating sugar for ECA. We generated evidence suggesting that *E. coli* K-12 redirects its UndPP-GlcNAc into ECA synthesis when OAg synthesis is disrupted, thereby alleviating the UndP sequestration stress. This was strongly supported with further experiments where inactivation of ECA synthesis renders *wzyB* essential in OAg producing *E. coli* K-12. In contrast, we found that disruptions of OAg synthesis in *S. flexneri* only marginally increased overall ECA production, suggesting that it was unable to redirect adequate UndPP-GlcNAc from the disrupted OAg synthesis pathways, resulting in cell death due to UndP sequestration.

Interestingly, we unexpectedly found only a limited *S. flexneri* cell population (30%) decorated their cell surface with ECA upon disruptions in OAg synthesis. Since all bacterial cells were grown from a single bacterial colony for all experimental repeats, it is tempting to suggest that the surface ECA decoration in *S. flexneri* may be phase variable. This is the first example of surface ECA being heterogeneously displayed on the surface of *S. flexneri* cells in the population, but it remains unclear what mechanism underpins this. Since polysaccharide synthesis including the ECA synthesis pathway, lacks a feedback control mechanism, whereby disruptions in late steps of biosynthesis cause UndP sequestration leading to cell death (26), it is therefore likely to be regulated at the committed biosynthesis step, i.e. the committed GT WecG and enzymes (WecB and WecC) responsible for the synthesis of its nucleotide sugar substrate. However, through sequence analysis between *S. flexneri* and *E. coli*, we were unable to identify any genetic regions that would potentially account for regulatory differences. Nevertheless, it may also be regulated through an additional molecular mechanism that is currently not known.

Polysaccharide polymerases were also reported to be essential for capsule synthesis in *Streptococcus pneumoniae* (27) and for ECA in *E. coli* K-12 (19, 28). However, ECA polymerase *wzyE* was found not essential in uropathogenic *E. coli* OAg-producing strains (2, 19), and here with an OAg restored *E. coli* K-12, highlighting the redirection of the common substrate UndPP-GlcNAc between ECA and OAg synthesis pathways as a mechanism to mitigate UndP sequestration stress. In addition, *wzyE* was found also essential in OAg producing *Salmonella* Typhimurium strains (19). This provides further supporting evidence, because the OAg for *Salmonella* Typhimurium initiates with galactose (29), different to the ECA initiating sugar GlcNAc, and is thereby unable to redirect UndP-GlcNAc into synthesis of OAg.

The diversification of the OAg RU structure presents significant challenges due to the high specificity of the enzymes involved in RU assembly and processing, such as GTs, translocases, and polymerases. When OAg structure is modified in the cytosolic face of IM, these existing enzymes may not recognise the altered UndPP-RU, leading to stalled synthesis. This disruption in the pathway can result in UndP sequestration and ultimately cause cell lysis, which potentially imposes evolutionary constraints for further diversification process. Interestingly, in contrast to *E. coli*, the diversification of *S. flexneri* OAg structures is primarily restricted to modifications introduced by bacterial phage elements, which encode enzymes modifying OAg structures on the periplasmic face of the inner membrane (30). This approach allows *S. flexneri* to bypass the specificity of the enzymes responsible for OAg synthesis. Our data suggest that, in addition to the pathogenic importance of *S. flexneri* OAg structures, a reduced capacity for ECA synthesis could pose an evolutionary constraint to its OAg diversification. Specifically, this limitation could impede the ability of *S. flexneri* to adapt its OAg structures efficiently through genetic recombination to acquire a new GT or other alterations leading to the evolution of a novel OAg structure, in that the limited tolerance to disrupted flows in OAg synthesis would quickly eliminate the recombinants before the acquisition of adaptive changes to evolve GT and flippase compatibility. We argue this may explain the restricted OAg structural diversity in *S. flexneri* compared to other *E. coli* lineages.

Most *E. coli, Shigella* and other Enterobacteriaceae have their OAg RU initiates with GlcNAc (or subsequently modified to GalNAc by epimerase on the lipid carrier in a reversible manner (9)) as their first sugar by the IT WecA (6) encoded in the ECA gene cluster, thereby sharing the first RU intermediate UndPP-GlcNAc with ECA RU synthesis. We propose a model in which sharing the initial sugar, GlcNAc, between OAg and ECA RUs allows for the redirection of UndPP-GlcNAc when one pathway encounters stalling or disruptions. This mechanism helps mitigate the stress caused by UndP sequestration (Fig 6A). Therefore, a ECA synthesis with high level of production would enhance the cell’s tolerance to UndP sequestration caused by disruptions in OAg synthesis. This increased production allows the cell to temporarily decorate its surface with ECA, which is crucial for maintaining cell survival while enabling further diversification of the new OAg (Fig. 6B). In contrast, a limited ECA production results in low tolerance to disruptions in OAg synthesis, leading to cell death and potentially impose constraints on the evolution process (Fig 6C).

**Fig 6.**
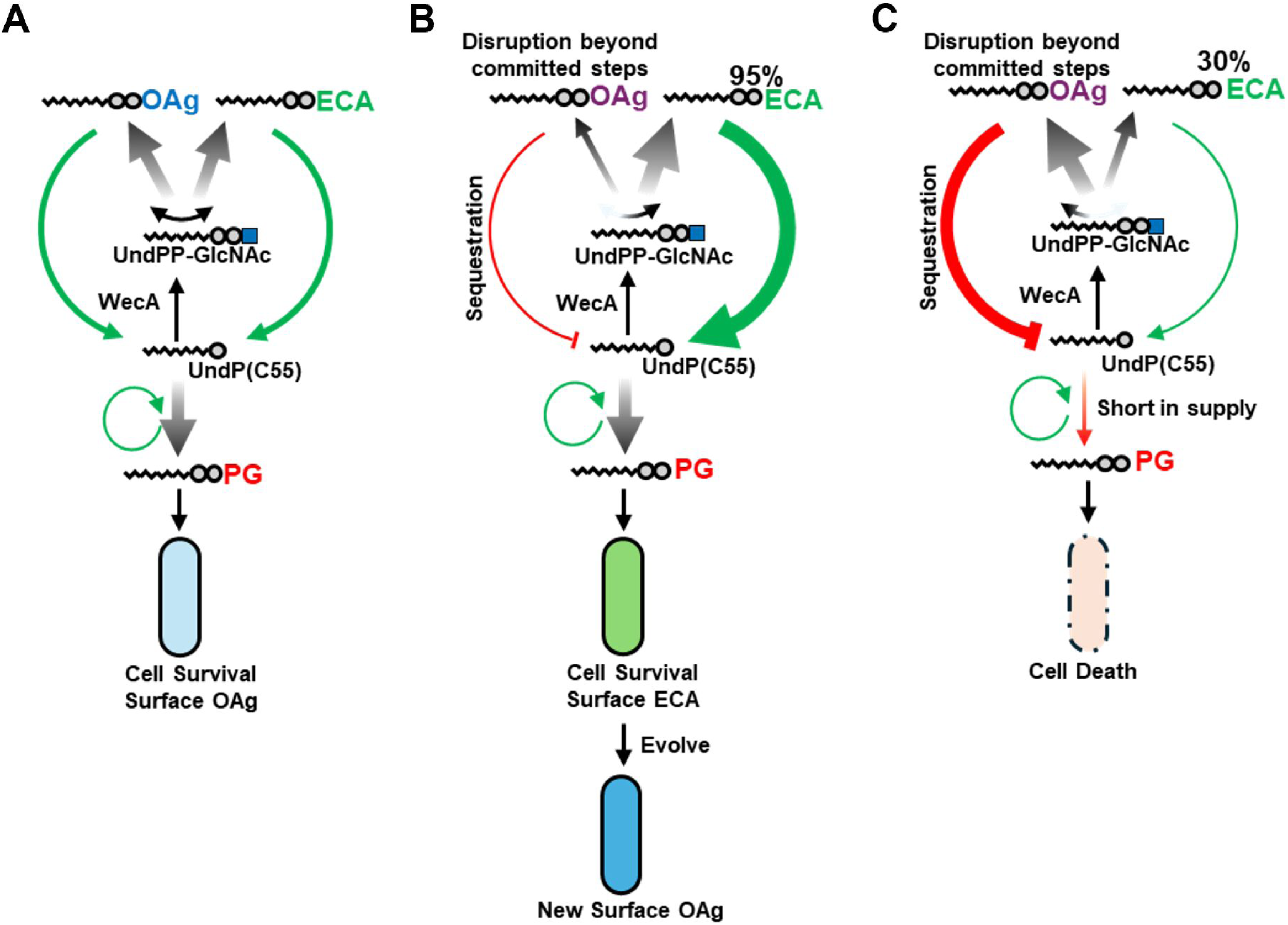
A model of the buffering mechanism that redirects UndP into ECA biosynthesis to maintain rapid UndP recycling during OAg pathway stalling or disruption. **A)** OAg and ECA RU assembly shares the initial substrate UndPP-GlcNAc which is catalysed by IT WecA to engage the lipid carrier UndP. The availability of UndP to PG biosynthesis is crucial for cell survival. When OAg biosynthesis is disrupted beyond the committed steps, UndPP-RU intermediates accumulate on the IM, locking UndP into the OAg biosynthesis pathway. **B)** This can be mitigated by redirecting UndPP-GlcNAc into a robust ECA biosynthesis pathway to maintain rapid UndP recycling rates. This cell survival supports further diversification of the new OAg. **C)** when the ECA pathway has limited biosynthesis capacity, disruptions in OAg biosynthesis may result in cell death halting further evolution process of OAg.

## Methods and Materials

### Bacterial strains and plasmids

Bacterial strains and plasmids used in this work are listed in Table 1. Single colonies grown on Lysogeny Broth (LB)-Lennox (31) agar plates were picked and grown overnight in LB at 37°C for all experiments. Where appropriate, LB media were supplemented with ampicillin (Amp, 100 µg/mL), kanamycin (Kan, 50 µg/mL), chloramphenicol (Chl, 25 µg/mL), anhydrotetracycline (AhTet 50 ng/mL), or arabinose (Ara, 10 mM).

**Table 1.**
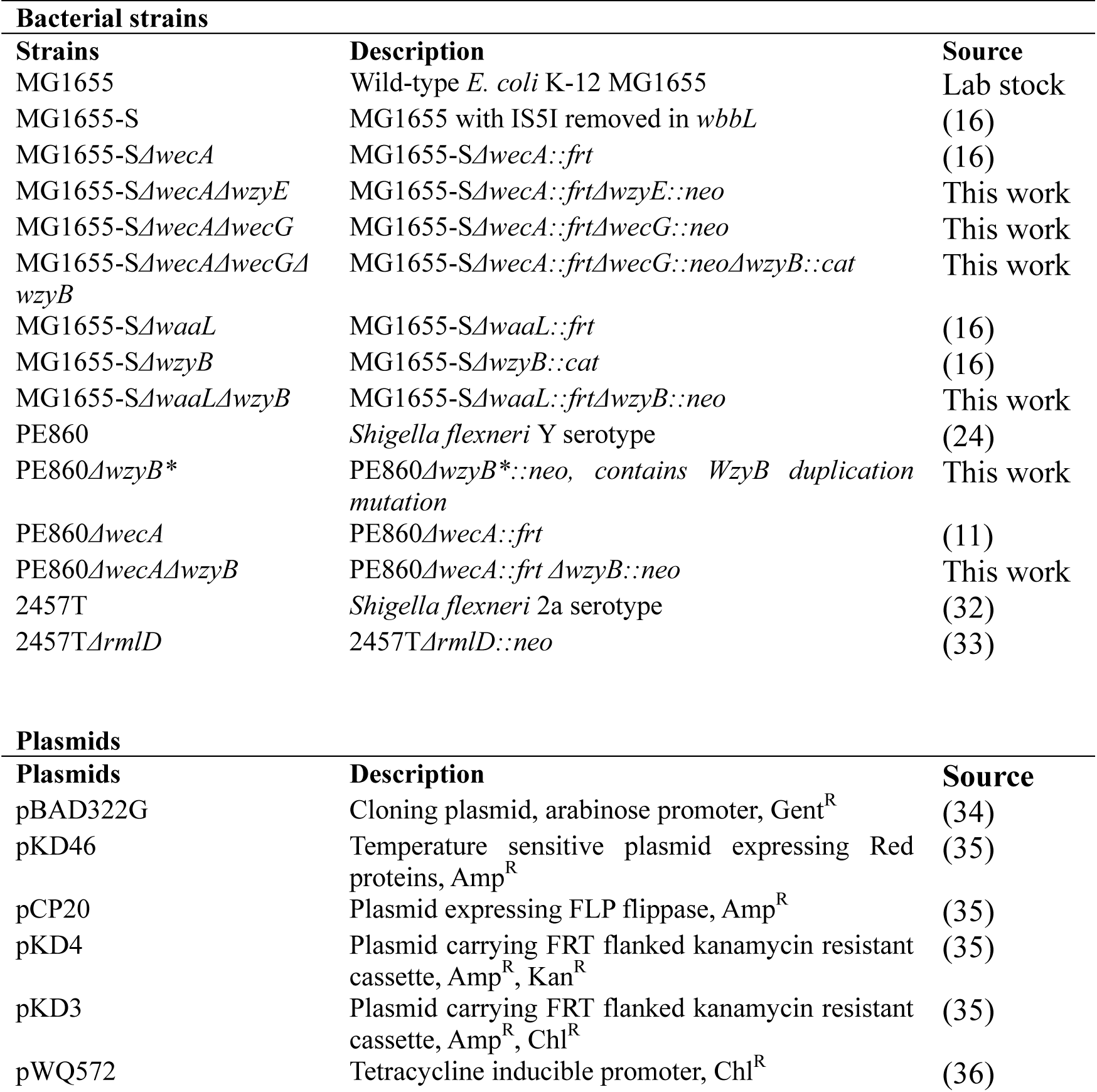
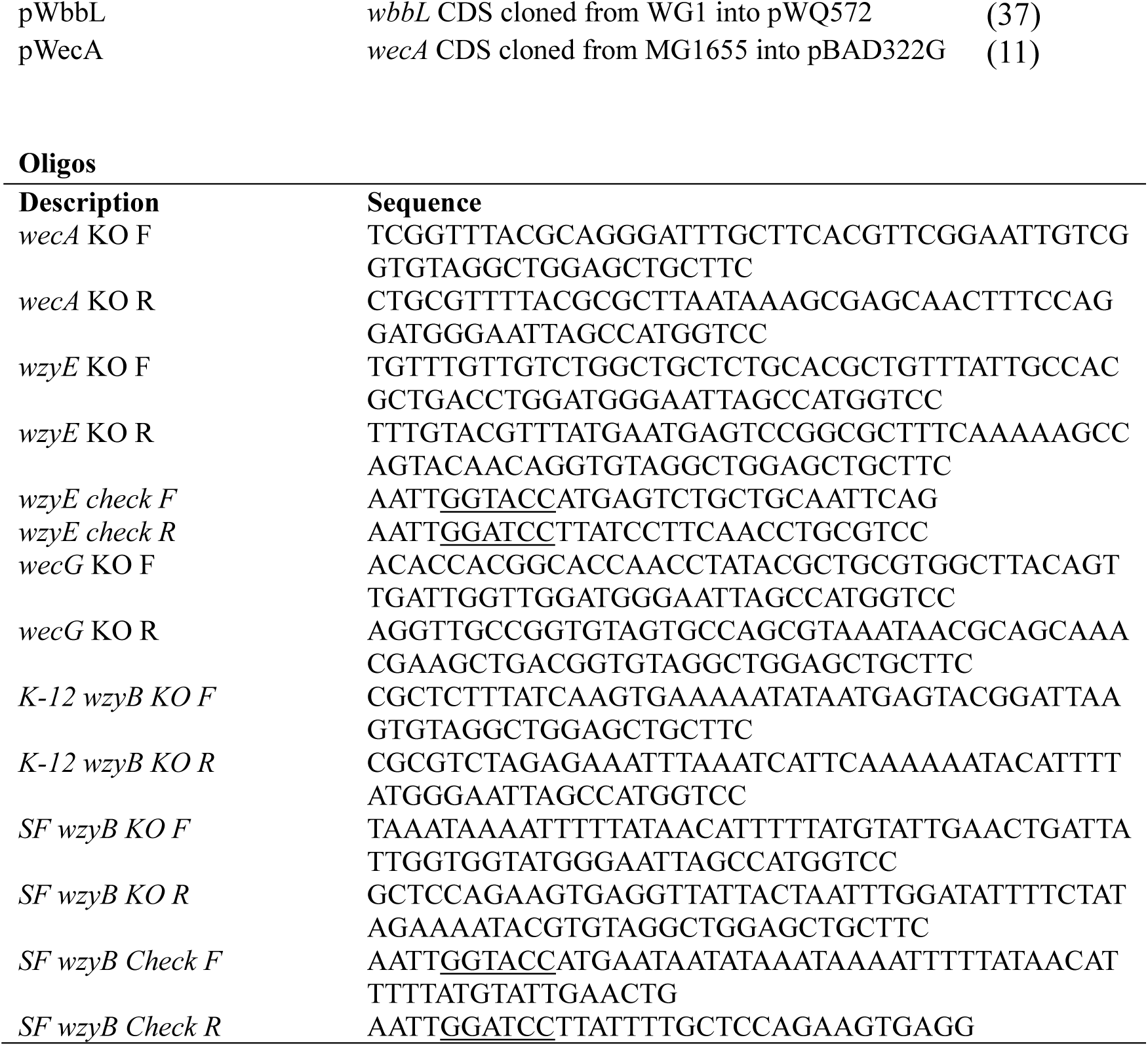
Strains, plasmids, and oligonucleotides.

### Bacterial mutagenesis via allelic exchange

Mutagenesis was performed as previously described (38) with adaptations (39). Bacterial strains with plasmid pKD46 were grown overnight in 10 mL LB at 30°C and diluted 1 in 100 into 10 mL LB. Lambda Red protein expression was induced with 50 mM L-arabinose when OD_600_ reached 0.3 and continued for 1 hour. Cells were then centrifuged, washed with ice-cold water, and resuspended in 100 µL of 10% ice-cold glycerol for electroporation. The *cat* or *neo* gene was PCR-amplified from pKD3 or pKD4, respectively, using primers with 40-50 bp of homologous sequences (Table 1). The purified PCR product (1.5 µg) was introduced into electrocompetent cells by electroporation. After recovery in 3 mL LB at 37°C for 2 hours, cells were plated on LB agar with Chl or Kan and incubated at 37°C for 16 hours. Mutants were then confirmed by PCR screening.

### Bacitracin sensitivity assay

Bacterial survival spotting assays were conducted as previously described (16). Overnight bacterial cultures were adjusted to an OD_600_ of 1.0, then serially diluted 10-fold to 10^−7^ in fresh LB media. A 4 µL aliquot of each dilution was spotted onto LB agar plates, with or without 1 mg/ml bacitracin (Sigma, B0125).

### Bacterial growth kinetic assay

Bacterial growth kinetics were recorded as described previously (16). Overnight bacterial cultures were diluted 1:200 in fresh LB media, with or without 10 mM arabinose and/or 50 ng/mL anhydrotetracycline, in a 96-well plate. The plate was incubated at 37°C with aeration, and OD_600_ was measured every 10 minutes for 18 hours using a CLARIOstar plate reader (BMG, Australia).

### Bacterial cell lysis assay

To measure bacterial cell lysis, WecA and WbbL were induced in bacterial cells grown to an OD_600_ of 0.8-1 and incubated for an additional 30 minutes at 37 °C. Cultures showing reduced turbidity due to cell lysis were imaged. The supernatants were then collected by centrifugation at 20,000 × g and mixed with 5 µg/mL Ethidium bromide (EtBr, BioRad). Fluorescence was measured using a CLARIOstar plate reader (BMG, Australia) with excitation at 525 nm and emission at 615 nm.

### Western immunoblotting

For western immunoblotting of ECA, bacterial cells (5◊10^8^) were harvested via centrifugation, and lysed in 50 µl of SDS sample buffer. Samples were then treated with 20 µg/ml proteinase K (NEB, #P8107S) overnight at 60 °C for 18 hours. Samples were then separated by SDS-Tris/Glycine gel (BioRad, #4568095) electrophoresis and subsequently transferred onto nitrocellulose membrane and detected with rabbit polyclonal anti-ECA antibodies (24). For loading control, lysed whole cell bacterial samples without proteinase K treatment were separated by SDS-PAGE and subsequently immunoblotted with anti-SurA antibodies (gifted by Carol Gross, University of California).

### Surface ECA immunostaining

For ECA surface labeling, bacteria (10^8^ cells) in mid-exponential phase were harvested by centrifugation (16,000 × g, 1 min), fixed with 3.7% (wt/vol) formaldehyde in PBS for 20 minutes at room temperature, and then washed with PBS. A 5 µL suspension of fixed bacteria was centrifuged onto coverslips precoated with 0.01% (wt/vol) poly-L-lysine (Sigma) in a 24-well tray (16,000 × g, 1 min). Coverslips were incubated sequentially with rabbit anti-ECA pAbs (1:100), followed by anti-rabbit Alexa Fluor 488 (Invitrogen, 1:100) in PBS with 10% (vol/vol) FBS (Gibco), with PBS washes in between. Coverslips were mounted with ProLong Diamond Antifade Mountant (Invitrogen) and imaged using a ZEISS Axio Vert.A1 microscope. The experiment was repeated once, and the percentage of ECA-positive bacteria was quantified using three micrographs.

## Acknowledgement

This work is funded by an Early Career Research Ideas Grant from the Faculty of Health, Queensland University of Technology (Australia) to JQ, and in part by an Australian Research Council project grant (DP210101317), the Max Planck Queensland Centre on the Materials Science of Extracellular Matrices to MT, The Ian

Potter Foundation sponsored the CLARIOStar high-performance microplate reader (BMG, Australia) and epifluorescence microscope. The funders had no role in study design, data collection and analysis, decision to publish, or preparation of the manuscript.

## Author contributions

JQ conceptualised the project; JQ and YH contributed to experimental design; JQ conducted experiments, and contributed to data collection, analysis and interpretation; JQ and MT supervised the study and obtained the funding. RM contributed to the generation of experimental materials. JQ wrote the manuscript, and all authors edited the manuscript.

## Data availability

All data generated or analysed during this study were included in this article.

## References

1. Chaput C, Spindler E, Gill RT, Zychlinsky A. 2013. O-antigen protects gram-negative bacteria from histone killing. PLoS One 8:e71097.

2. Phan MD, Peters KM, Sarkar S, Lukowski SW, Allsopp LP, Gomes Moriel D, Achard ME, Totsika M, Marshall VM, Upton M, Beatson SA, Schembri MA. 2013. The serum resistome of a globally disseminated multidrug resistant uropathogenic Escherichia coli clone. PLoS Genet 9:e1003834.

3. Smith HW. 1975. Survival of orally administered E. coli K 12 in alimentary tract of man. Nature 255:500–2.

4. Day CJ, Tran EN, Semchenko EA, Tram G, Hartley-Tassell LE, Ng PS, King RM, Ulanovsky R, McAtamney S, Apicella MA, Tiralongo J, Morona R, Korolik V, Jennings MP. 2015. Glycan:glycan interactions: High affinity biomolecular interactions that can mediate binding of pathogenic bacteria to host cells. Proc Natl Acad Sci U S A 112:E7266–75.

5. West NP, Sansonetti P, Mounier J, Exley RM, Parsot C, Guadagnini S, Prevost MC, Prochnicka-Chalufour A, Delepierre M, Tanguy M, Tang CM. 2005. Optimization of virulence functions through glucosylation of Shigella LPS. Science 307:1313–7.

6. Liu B, Furevi A, Perepelov AV, Guo X, Cao H, Wang Q, Reeves PR, Knirel YA, Wang L, Widmalm G. 2020. Structure and genetics of Escherichia coli O antigens. FEMS Microbiol Rev 44:655–683.

7. Kauffmann F. 1947. The serology of the coli group. J Immunol 57:71–100.

8. Barreteau H, Magnet S, El Ghachi M, Touze T, Arthur M, Mengin-Lecreulx D, Blanot D. 2009. Quantitative high-performance liquid chromatography analysis of the pool levels of undecaprenyl phosphate and its derivatives in bacterial membranes. J Chromatogr B Analyt Technol Biomed Life Sci 877:213–20.

9. Rush JS, Alaimo C, Robbiani R, Wacker M, Waechter CJ. 2010. A novel epimerase that converts GlcNAc-P-P-undecaprenol to GalNAc-P-P-undecaprenol in Escherichia coli O157. J Biol Chem 285:1671–80.

10. Liu D, Reeves PR. 1994. Escherichia coli K12 regains its O antigen. Microbiology (Reading) 140 (Pt 1):49–57.

11. Qin J, Hong Y, Totsika M. 2024. Determining glycosyltransferase functional order via lethality due to accumulated O-antigen intermediates, exemplified with Shigella flexneri O-antigen biosynthesis. Appl Environ Microbiol 90:e0220323.

12. Hong Y, Cunneen MM, Reeves PR. 2012. The Wzx translocases for Salmonella enterica O-antigen processing have unexpected serotype specificity. Mol Microbiol 84:620–30.

13. Hong Y, Morcilla VA, Liu MA, Russell ELM, Reeves PR. 2015. Three Wzy polymerases are specific for particular forms of an internal linkage in otherwise identical O units. Microbiology (Reading) 161:1639–1647.

14. Wang L, Reeves PR. 1998. Organization of Escherichia coli O157 O antigen gene cluster and identification of its specific genes. Infect Immun 66:3545–51.

15. Liu D, Cole RA, Reeves PR. 1996. An O-antigen processing function for Wzx (RfbX): a promising candidate for O-unit flippase. J Bacteriol 178:2102–7.

16. Qin J, Hong Y, Morona R, Totsika M. 2023. O antigen biogenesis sensitises Escherichia coli K-12 to bile salts, providing a plausible explanation for its evolutionary loss. PLoS Genet 19:e1010996.

17. Jorgenson MA, Young KD. 2016. Interrupting Biosynthesis of O Antigen or the Lipopolysaccharide Core Produces Morphological Defects in Escherichia coli by Sequestering Undecaprenyl Phosphate. J Bacteriol 198:3070–3079.

18. Freed NE, Bumann D, Silander OK. 2016. Combining Shigella Tn-seq data with gold-standard E. coli gene deletion data suggests rare transitions between essential and non-essential gene functionality. BMC Microbiol 16:203.

19. Ghomi FA, Jung JJ, Langridge GC, Cain AK, Boinett CJ, Abd El Ghany M, Pickard DJ, Kingsley RA, Thomson NR, Parkhill J, Gardner PP, Barquist L. 2024. High-throughput transposon mutagenesis in the family Enterobacteriaceae reveals core essential genes and rapid turnover of essentiality. mBio doi:10.1128/mbio.01798-24:e0179824.

20. Morona R, Mavris M, Fallarino A, Manning PA. 1994. Characterization of the rfc region of *Shigella flexneri*. J Bacteriol 176:733–47.

21. Nath P, Tran EN, Morona R. 2015. Mutational analysis of the Shigella flexneri O-antigen polymerase Wzy: identification of Wzz-dependent Wzy mutants. J Bacteriol 197:108–19.

22. Tran EN, Papadopoulos M, Morona R. 2014. Relationship between O-antigen chain length and resistance to colicin E2 in *Shigella flexneri*. Microbiology 160:589–601.

23. Ascari A, Tran ENH, Eijkelkamp BA, Morona R. 2022. Identification of the Shigella flexneri Wzy Domain Modulating Wzz(pHS-2) Interaction and Detection of the Wzy/Wzz/Oag Complex. J Bacteriol 204:e0022422.

24. Maczuga N, Tran ENH, Qin J, Morona R. 2022. Interdependence of Shigella flexneri O Antigen and Enterobacterial Common Antigen Biosynthetic Pathways. J Bacteriol 204:e0054621.

25. Yuasa R, Levinthal M, Nikaido H. 1969. Biosynthesis of cell wall lipopolysaccharide in mutants of Salmonella. V. A mutant of Salmonella typhimurium defective in the synthesis of cytidine diphosphoabequose. J Bacteriol 100:433–44.

26. Eade CR, Wallen TW, Gates CE, Oliverio CL, Scarbrough BA, Reid AJ, Jorgenson MA, Young KD, Troutman JM. 2021. Making the Enterobacterial Common Antigen Glycan and Measuring Its Substrate Sequestration. ACS Chem Biol 16:691–700.

27. Xayarath B, Yother J. 2007. Mutations blocking side chain assembly, polymerization, or transport of a Wzy-dependent Streptococcus pneumoniae capsule are lethal in the absence of suppressor mutations and can affect polymer transfer to the cell wall. J Bacteriol 189:3369–81.

28. Gerdes SY, Scholle MD, Campbell JW, Balazsi G, Ravasz E, Daugherty MD, Somera AL, Kyrpides NC, Anderson I, Gelfand MS, Bhattacharya A, Kapatral V, D’Souza M, Baev MV, Grechkin Y, Mseeh F, Fonstein MY, Overbeek R, Barabasi AL, Oltvai ZN, Osterman AL. 2003. Experimental determination and system level analysis of essential genes in Escherichia coli MG1655. J Bacteriol 185:5673–84.

29. Wang L, Andrianopoulos K, Liu D, Popoff MY, Reeves PR. 2002. Extensive variation in the O-antigen gene cluster within one Salmonella enterica serogroup reveals an unexpected complex history. J Bacteriol 184:1669–77.

30. Sadredinamin M, Yazdansetad S, Alebouyeh M, Yazdi MMK, Ghalavand Z. 2023. Shigella Flexneri Serotypes: O-antigen Structure, Serotype Conversion, and Serotyping Methods. Oman Med J 38:e522.

31. Lennox ES. 1955. Transduction of linked genetic characters of the host by bacteriophage P1. Virology 1:190–206.

32. Qin J, Doyle MT, Tran ENH, Morona R. 2020. The virulence domain of Shigella IcsA contains a subregion with specific host cell adhesion function. PLoS One 15:e0227425.

33. Van den Bosch L, Morona R. 2003. The actin-based motility defect of a *Shigella flexneri* rmlD rough LPS mutant is not due to loss of IcsA polarity. Microb Pathog 35:11–8.

34. Cronan JE. 2006. A family of arabinose-inducible Escherichia coli expression vectors having pBR322 copy control. Plasmid 55:152–7.

35. Datsenko KA, Wanner BL. 2000. One-step inactivation of chromosomal genes in Escherichia coli K-12 using PCR products. Proceedings of the National Academy of Sciences of the United States of America 97:6640–6645.

36. Larue K, Ford RC, Willis LM, Whitfield C. 2011. Functional and structural characterization of polysaccharide co-polymerase proteins required for polymer export in ATP-binding cassette transporter-dependent capsule biosynthesis pathways. J Biol Chem 286:16658–68.

37. Hong Y, Reeves PR. 2014. Diversity of o-antigen repeat unit structures can account for the substantial sequence variation of wzx translocases. J Bacteriol 196:1713–22.

38. Datsenko KA, Wanner BL. 2000. One-step inactivation of chromosomal genes in Escherichia coli K-12 using PCR products. Proc Natl Acad Sci U S A 97:6640–5.

39. Qin J, Hong Y, Pullela K, Morona R, Henderson IR, Totsika M. 2022. A method for increasing electroporation competence of Gram-negative clinical isolates by polymyxin B nonapeptide. Sci Rep 12:11629.

